# Real-time detection of bursts in neuronal cultures using a Neuromorphic Auditory Sensor and Spiking Neural Networks

**DOI:** 10.1101/2020.05.20.105593

**Authors:** Juan P. Dominguez-Morales, Stefano Buccelli, Daniel Gutierrez-Galan, Ilaria Colombi, Angel Jimenez-Fernandez, Michela Chiappalone

**Affiliations:** Robotics and Technology of Computers Lab. Universidad de Sevilla, Spain; Rehab Technologies IIT-INAIL Lab, Istituto Italiano di Tecnologia, Via Morego 30, 16163 Genova, Italy; Department of Neuroscience and Brain Technologies, Istituto Italiano di Tecnologia, Via Morego 30, 16163 Genova, Italy

**Keywords:** SpiNNaker, Spiking Neural Networks, Neuromorphic Hardware, Brain Signals Processing, Burst detection

## Abstract

The correct identification of burst events is crucial in many scenarios, ranging from basic neuroscience to biomedical applications. However, none of the burst detection methods that can be found in the literature have been widely adopted for this task. As an alternative to conventional techniques, a novel neuromorphic approach for real-time burst detection is proposed and tested on acquisitions from in vitro cultures. The system consists of a Neuromorphic Auditory Sensor, which converts the input signal obtained from electrophysiological recordings into spikes and decomposes them into different frequency bands. The output of the sensor is sent to a trained spiking neural network implemented on a SpiNNaker board that discerns between bursting and non-bursting activity. This data-driven approach was compared with 8 different conventional spike-based methods, addressing some of their drawbacks, such as being able to detect both high and low frequency events and working in an online manner. Similar results in terms of number of detected events, mean burst duration and correlation as current state-of-the-art approaches were obtained with the proposed system, also benefiting from its lower power consumption and computational latency. Therefore, our neuromorphic-based burst detection paves the road to future implementations for neuroprosthetic applications.

## 1. Introduction

The analysis of neural signals in real-time is a hot topic in both neuroscience and neuroengineering fields. Current applied research is focusing on the development of closed-loop devices [1, 2] for treating a wide variety of pathologies such as epilepsy, Parkinson’s disease, chronic pain, stroke and mood disorders [3, 4, 5, 6, 7, 8]. Those systems typically rely on the real-time recognition of specific patterns of activity, which can be used to trigger an electrical therapy [9]. In order to develop and test appropriate algorithms for real-time pattern detection, simple experimental models could be used, capable to exhibit peculiar events of activity typical of higher and more complex organisms [10].

In this regard, neuronal cultures plated over Micro Electrode Arrays (MEAs) have been proposed as an excellent test bed for studying relevant electrophysiological events [11, 12]. One peculiar feature of neural network activity resembled by mature neural cultures in vitro is the presence of synchronized and packed spiking activity known as bursts [13, 14, 15], distributed throughout the network. Given the wide spectrum of spontaneous bursting activity patterns, several techniques for detecting those events have been developed. However, none of them are commonly recognized as the best for all criteria [16]. Typically, burst detection algorithms are based on an initial processing of spike detection, which lacks on a definitive method for all neurons and noise conditions [17]. For burst-like epileptic discharges, other raw-based methods have been used in the literature and in the clinical practice [18, 19]. In this paper, a different approach to the problem of burst detection is considered. Instead of using spike detection as a first step of the process, we used wide band signals and exploited a neuromorphic signal processing system to detect bursts. It is worth highlighting that neuromorphic-based algorithms have become an alternative to traditional classification systems due to the interest on bio-inspired processing techniques and the increased demand for low-power and low-latency platforms [20, 21].

The first step of the proposed processing system involves the use of a Neuromorphic Auditory Sensor (NAS) [22] implemented on a Field Programmable Gate Array (FPGA). NAS was designed to work with spikes to mimic the information coding of a biological cochlea by means of a frequency decomposition. Such decomposition carries information both from high frequency (i.e., the frequency range in which biological spikes can be recorded) and low frequency components (i.e. local field potential, LFP). After this innovative processing stage, the resulting spikes constitute the input for a Spiking Neural Network (SNN) implemented by means of the SpiNNaker platform, which is a massively-parallel multicore computing system designed for modeling very large SNNs in real time [23].

For the purpose of this study, a 2-layer SNN was designed to perform a classification based on the decomposition obtained from the NAS. The SNN was trained on a subset of the recorded dataset by means of the Spike-Timing-Dependent Plasticity (STDP) learning algorithm [24, 25]. Then, the trained setup was used to infer a decision over the non-trained dataset, and the results were compared with state-of-the-art algorithms for burst detection.

The paper is structured as follows: section 2 presents the materials and methods used, describing both biological experiments (2.1), neuromorphic-based processing (2.2), spike-based processing (2.3), raw-based processing (2.4) and statistical analysis (2.5). Finally, the results of this work are presented in section 3, along with a discussion and future perspectives (section 4).

## 2. Materials and Methods

The electrophysiological signals recorded from the MEA setup (see Fig. 1 and section 2.1) were processed in different ways. Specifically, we performed neuromorphic processing, a conventional spike-based processing and a raw-based processing as depicted in Fig. 1. The neuromorphic process includes a software cubic spline interpolation of the signal (section 2.2.1), resulting in 48kHz raw data (i.e., the audio signal). This signal is analyzed by the Neuromorphic Auditory Sensor (NAS), which decomposes it in 128 spike-trains (events) (section 2.2.2). The output of the NAS (i.e., the spike events obtained upon the frequency decomposition) serves as input for the SNN model implemented on the SpiNNaker board (section 2.2.3), which implements the neuromorphic burst detection. The first step of the conventional process is a software high-pass filter at 300Hz, followed by a spike detection module (see section 2.3.2) and finally the resulting burst train is compared with the other approaches. The raw-based process (see section 2.4) includes binning the signal a in 50ms-long time window and performing a raw-based burst detection.

**Figure 1:**
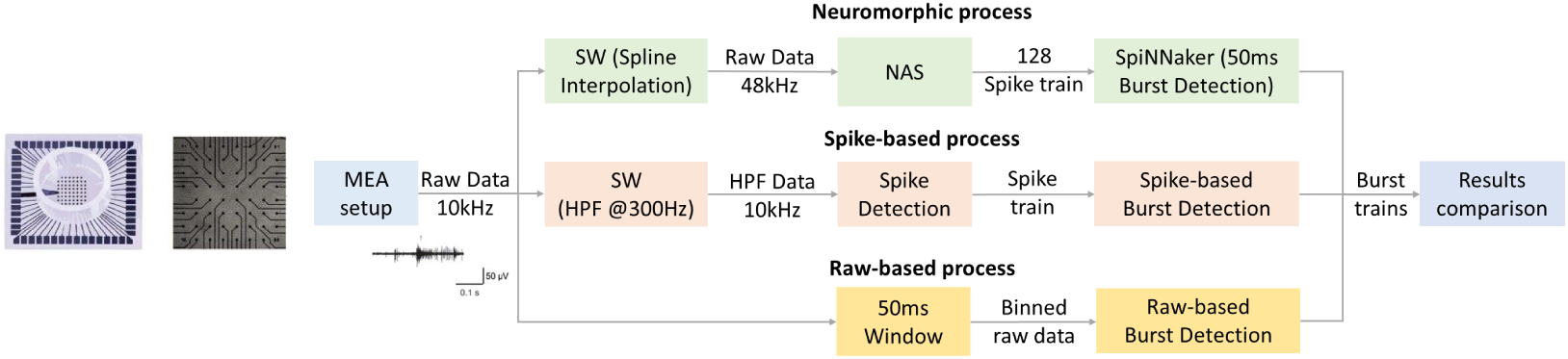
Overview of the methodological approach. 60-channel-MEA (Multichannel System, MCS, Reutlingen, Germany) with the standard electrode layout (8 × 8) plated with cortical cultures. Schematic description of the different burst detection approaches. The signal acquired from the MEA setup (at 10kHz) is processed with different approaches. The spike-based process includes a software high-pass filter at 300Hz, followed by a spike detection and a spike-based burst detection. The raw-based process includes binning the signal in a 50ms-long time window and performing a raw-based burst detection. The neuromorphic process includes a software cubic spline interpolation of the signal, resulting in 48kHz raw data. This signal is analyzed by the Neuromorphic Auditory Sensor (NAS), which decomposes the signal in 128 spike-trains (addresses). The 128 addresses are analyzed by the neuromorphic SpiNNaker burst detection module in 50ms time windows.

### 2.1. Biological experiments

#### 2.1.1. Cell cultures

Dissociated neuronal cultures were prepared from neocortex of embryonic rats at gestational day 18 (pregnant Sprague-Dawley female rats delivered by Charles River Laboratories). The procedures for preparing neuronal cultures are described in detail in previous studies [26, 12].

After enzymatic digestion in trypsin solution 0.125% (30 min at 37^*°*^C) and mechanical dissociation, the resulting tissue was re-suspended in Neurobasal medium supplemented with 2% B27, 1% Glutamax-1, 1% Pen-Strep solution, and 10% FBS (Invitrogen, Carlsbad, CA, USA), at the final concentration of 1500 cells/*µ*l. Cells were afterward plated onto standard 60-channel Micro Electrode Arrays (MEAs) (Multichannel Systems, MCS, Reutlingen, Germany) previously coated with poly-D-lysine and laminin (final density around 1200 cells/mm^2^) and maintained with 1 ml of nutrient medium in a humidified incubator (5% CO2 and 95% air at 37^*°*^C). Half of the medium was replaced weekly.

#### 2.1.2. Micro-electrode array recordings

Planar microelectrodes were arranged in an 8 × 8 layout, excluding corners and one reference electrode, for a total of 59 TiN/SiN planar round recording electrodes (30*µ*m diameter; 200*µ*m center-to-center inter electrode distance), which can be seen in Fig. 1. One recording electrode was replaced with a bigger ground electrode. The activity was recorded by means of the MEA60 System (Multichannel Systems, MCS, Reutlingen, Germany). The signal from each electrode was sampled at 10 kHz and amplified with a band-width of 1 Hz–3 kHz. Each recorded electrode was acquired through the data acquisition card and on-line monitored through MC Rack software (Multi-channel Systems, MCS, Reutlingen, Germany). To reduce thermal stress of the cells during the experiment, MEAs were kept at 37^*°*^C by means of a controlled thermostat and covered by polydimethylsiloxane (PDMS) caps to avoid evaporation and prevent changes in osmolarity.

#### 2.1.3. Experimental protocols and dataset

We recorded the electrophysiological activity from eight cortical cultures plated over MEA, with an average age of 31 ± 2.1 DIV (Days In Vitro) to have a stable bursting activity [27]. The spontaneous activity was monitored and recorded for 5 minutes, after a period of 30 minutes rest outside the incubator into the experimental set-up, to let the culture adapt to the new environment and reach a stable level of activity [28]. We split the dataset as follows: one electrode from the first culture was used as training set and 14 electrodes belonging to the other seven cultures (2 for each of them) were used as test set. The choice of the two channels was made to obtain the largest variability in terms of signal-to-noise ratio among the bursting channels. Regarding the training set, we performed a visual inspection to split it in “bursting” and “non-bursting”.

### 2.2. Neuromorphic-based processing

#### 2.2.1. Preprocessing

In order to be used by the NAS [22], the electrophysiological signals were converted to audio signals. A cubic spline interpolation was used to reach a sampling frequency of 48 kHz in order to match the sampling frequency of the mixer that was used to play the audio signal back to the NAS.

#### 2.2.2. Neuromorphic Auditory Sensor (NAS)

A NAS [22] is a neuromorphic digital audio sensor implemented on FPGA inspired by Lyon’s model of the biological cochlea [29]. This neuromorphic sensor is able to process excitatory audio signals using Spike Signal Processing (SSP) techniques [30], decomposing incoming audio in its frequency components, and providing this information as a stream of events using the Address-Event Representation (AER) [31].

This decomposition is carried out by a series of cascade-connected stages, modeling the basilar membrane of the biological cochlea, which decomposes the input audio signal into different frequency bands (also called channels). The entire system has been described in a previous publication [22]. For the sake of clarity, we here report the main components and their functions (see Fig. 2). Panel A of Fig. 2 shows a picture of the hardware setup used, which included a 64-channel monaural NAS implemented on an AER-Node board (1), an I2S A/D converter audio input interface for the board (2), a 3.3V-to-1.8V adapter PCB (3), a 4-chip SpiNNaker board (4), and the ethernet interface to communicate the SpiNNaker board with the PC (5).

**Figure 2:**
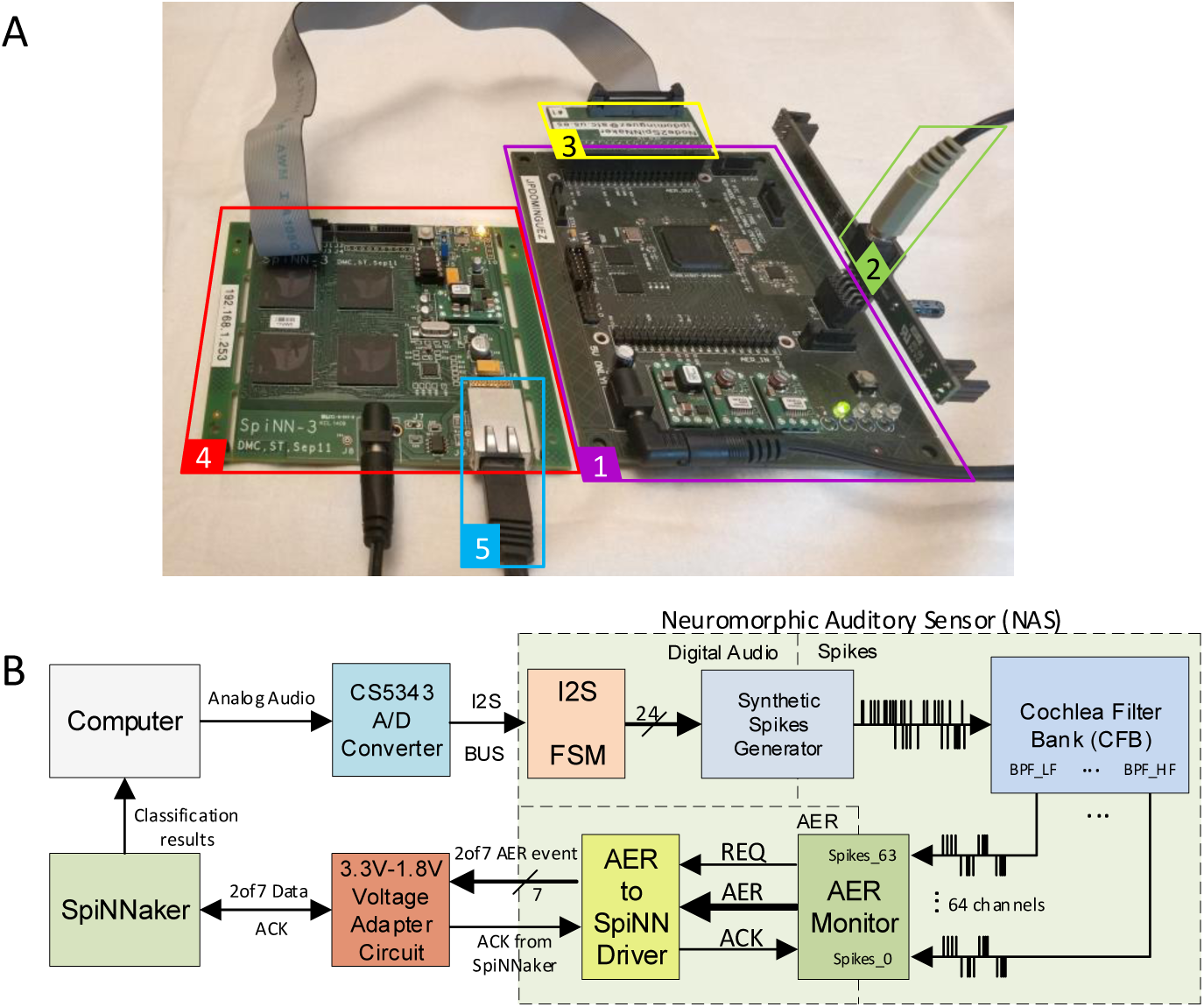
The neuromorphic system. A) Picture depicting the entire neuromorphic setup: 1) NAS implemented on the AER-Node board, 2) audio input, 3) 3.3V-to-1.8V adapter PCB, 4) SpiNNaker board, 5) SpiNNaker-to-PC ethernet interface. B) Block diagram of the setup used for the neuromorphic system, where *I2S* stands for Integrated Interchip Sound; *FSM*, for Finite State Machine; *REQ*, for Request; and *ACK*, for Acknowledge.

To digitize audio signals, a CS5344 Analog-to-Digital Converter (A/D) with a resolution of 24 bits and a sample rate of 96 KSamples/s is used (Fig. 2 panel B: A/D converter and Integrated Interchip Sound (I2S) Finite State Machine (FSM)). The digital audio signal is then converted to spikes using the spike generator presented in [32] (Fig. 2 panel B: Synthetic Spikes Generator), and used as input to the NAS filters (Fig. 2 panel B: Cochlea Filter Bank (CFB)).

Output spikes from the CFB are connected to an AER monitor [33], giving a unique address (which corresponds to the NAS channel) to the generated spikes following the AER protocol, and propagating them using an asynchronous AER bus.

A 64-channel monaural NAS implemented on an AER-Node board (shown in Fig. 2 panel A with number 1), which is based on a Spartan-6 FPGA (shown in Fig 2 panel B as NAS), was used in this work to process the audio signals. The NAS, which has 64 channels, generates spikes with addresses that range from 0 to 127, since each channel has two different spike trains that correspond to the positive and negative part of the signal, respectively. The output spikes from the sensor are first converted to the 2-of-7 protocol that SpiNNaker needs using an AER to SpiNN driver and then sent to a 4-chip SpiNNaker machine (called SpiNN-3) using a NAS-to-SpiNNaker PCB interface (Fig. 2 panel B: 3.3V-1.8V Voltage Adapter Circuit) in order to classify the activity within the signal between *non-bursting* and *bursting activity*. Thus, the NAS was used as input to the SNN.

#### 2.2.3. Spiking Neural Network Architecture (SpiNNaker)

As mentioned in the previous section, in this work a 4-chip SpiNNaker board (SpiNN-3) [34] was used, which can be seen in Panel A Fig. 2). Each chip consists of eighteen 200 MHz general-purpose low-power ARM968 cores, for a total of 72 ARM processor cores. A 100 Mbps Ethernet connection is used as control and I/O interface between the computer and the SpiNNaker board. This platform has been used in previous works by the authors for many different tasks, such as audio classification [35, 36], speech recognition [37], pattern recognition [38] and for building Central Pattern Generators (CPGs) [39, 40], proving its robustness and versatility.

SpiNN-3 also has two SpiNNaker links to connect other devices such as FPGAs or neuromorphic sensors like retinas and cochleas [41, 42]. In this work, we used one of these links to connect the NAS in order to feed the SNN implemented on SpiNNaker directly with the output information from the NAS, thus allowing real-time classification.

#### 2.2.4. Training the network with STDP

STDP [43, 44, 24, 45] is a biological process that is able to adjust the strength (weights) of the connections between neurons based on the relative timing of the output of a particular neuron and the input spiking activity. STDP has been implemented in SNN simulators like Brian [46] or NEST [47], and even in SPyNNaker [48]^1^, which is the software layer running in SpiNNaker machines. It is the most common training mechanism for spike-based networks, and it allows an easy and relatively fast online training phase of the network [25].

In this work, the network architecture shown in Panel A of Fig. 3 was used to train the SNN. A teaching neuron was used to perform a supervised training step, forcing the output neuron to spike when a specific input pattern was injected, and thus, increasing and adjusting the weights of the projections between the input and the output neuron that corresponds to that input pattern. After training, the teaching neuron was removed and STDP connections with the trained weights were kept in order to test the system. These weights can be seen in Panel B of Fig. 3.

**Figure 3:**
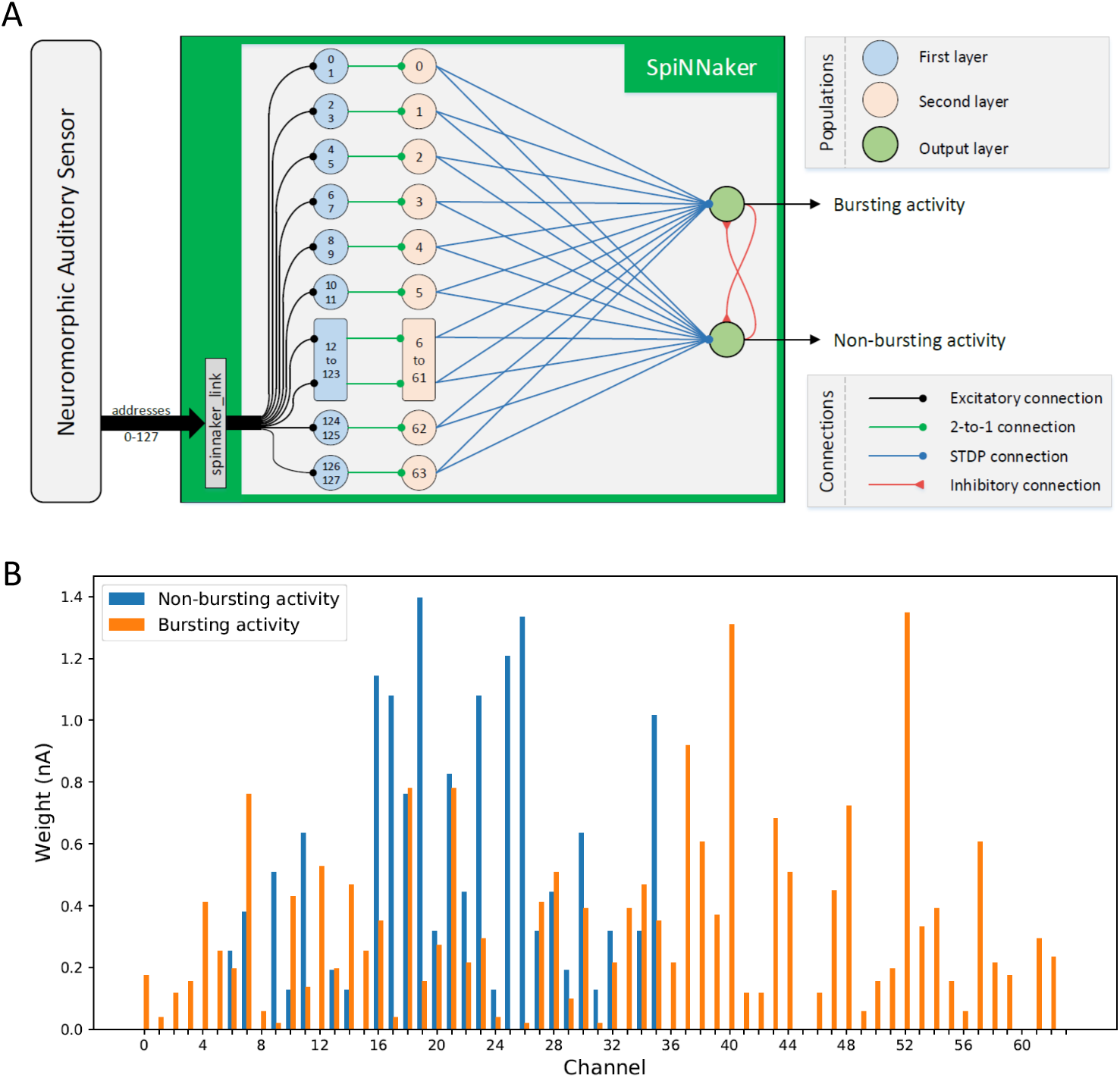
SNN learning process. A) Spiking Neural Network architecture. B) Weights obtained in the STDP training process. Blue values correspond to the weights (in nanoamperes) of the projections between the second layer and *non-bursting activity* class neuron from the output layer, while the orange values correspond to the weights from the neurons in the second layer to the *burst activity* neuron in the output layer. Higher IDs correspond to the information from lower frequencies.

For the training, the activity recorded from the selected bursting electrode (belonging to the first culture) was visually inspected and classified in bursting and non-bursting activity. We then concatenated the non-consecutive parts of the signal sharing the same class, obtaining 4 minutes and 41 seconds of non-bursting and 19 seconds of bursting activity (Fig. 8), which were used to train the SNN. For the testing, we used the trained SNN on 14 different recordings, two channels belonging to the remaining seven cultures.

#### 2.2.5. SNN-based burst detection

Once trained, the SNN was tested on different recordings. The output of the burst classification can be seen in real time; however, in order to compare this approach with others and measure its performance, the results were saved in a text file for further processing. The SpiNNaker script generated a text file containing the classification performed by the SNN according to a time window set at 50 ms. The output classes can be: “no signal” (when no audio is played), “bursting” and “non-bursting”.

A custom Matlab script counted the number of bursts that occurred in an audio recording based on the output results obtained from SpiNNaker. Moreover, the starting and ending time of each burst were stored for further analysis.

### 2.3. Spike-based processing

As anticipated at the beginning of the Materials and Methods section, we compared the NAS burst detection with other state-of-the-art methods as described in the following paragraphs.

#### 2.3.1. Preprocessing and spike detection

A conventional data analysis was performed offline by using a custom software developed in Matlab called SPYCODE [49], which collects a series of tools for processing multi-channel neural recordings.

An offline data filtering by means of a high-pass Butterworth filter with cut-off frequency at 300 Hz was performed in order to select only the Multi-Unit Activity (MUA) components of the signal, as reported in the literature [50, 51].

The Precise Timing Spike Detection (PTSD) algorithm [52] was used to discriminate spike events and to isolate them from noise by means of three parameters: (1) a differential threshold (DT), which was set independently for each channel and computed as 8 times the standard deviation (SD) of the noise of the signal; (2) a peak lifetime period (PLP) set to 2 ms; and (3) a refractory period set to 1 ms.

#### 2.3.2. Spike-based burst detection methods

Based on the work and open-source R code provided in [16], we performed a comparative analysis between 8 conventional, spike-based, burst detection methods and ours. We chose the following methods:

- CMA (cumulative moving average) [53]
- ISIrank threshold [54]
- PS (Poisson Surprise) [54]
- RS (Rank Surprise) [55]
- LogISI (Logarithmic Inter Spike Interval) [56]
- MI (Max Interval) [16]
- HSMM (Hidden Semi-Markov Model) [57]
- CH (Chiappalone, similar to max interval) [58]

Once the spike and burst detection procedures were performed, we extracted other parameters describing the electrophysiological patterns, such as the number of bursts and the average burst duration (ms). It is worth highlighting that all these methods are spike-based, meaning that they need a spike detection procedure to work. To further investigate this activity, a visual inspection (VI) on the high-pass filtered data was performed by an expert electrophysiologist. The VI trace was then used as our ground truth. All the parameters used for this study can be found in Table 1.

**Table 1:**
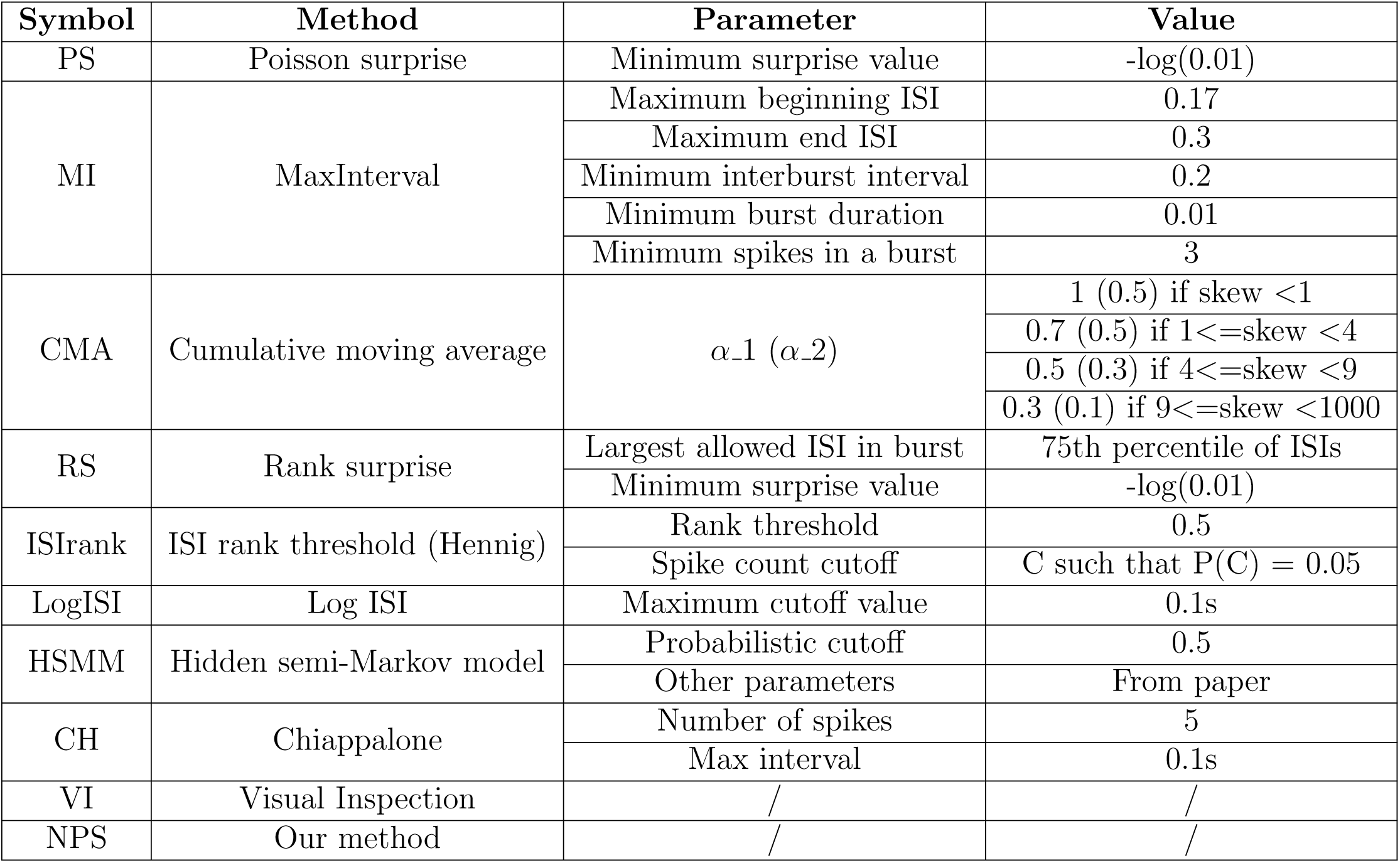
Burst detection parameters.

### 2.4. Raw-based processing

Besides the spike-based algorithms for burst detection, we compared NAS with other burst detectors designed to work on the raw data. We first split the raw data on 50ms time windows. Then, in order to discriminate between bursting and non bursting activity, we computed the following three different features for each time window:

- *max*: the maximum value in the time window;
- *peak peak*: the peak to peak difference in the time window;
- *len*: the signal length (defined as the sum of the absolute values of the first derivative of the signal).

A simple threshold was then used to separate bursting from non-bursting time windows. The best threshold, for each method was the one that maximized sensibility and specificity, and it was obtained as the point that minimized the distance from the top-left corner of the Receiver Operating Characteristic (ROC) curve [59]. We used the code provided by Víctor Martínez-Cagigal, ROC Curve (https://www.mathworks.com/matlabcentral/fileexchange/52442-roc-curve), from MATLAB Central File Exchange, retrieved April 01, 2020).

This threshold was the one which better discriminated the histograms of the two classes (i.e., bursting vs non-busting) with respect to the ground truth (see Fig. 9). Once we found the thresholds for the training set, we applied them to the test set recordings.

### 2.5. Statistical analysis

Given the non-normality of the distributions, we employed the Kruskal-Wallis test to compare the results obtained from different burst detection methods. Then, we employed the Bonferroni post-hoc analysis to investigate specific differences between methods. The statistical analyses were carried out using Matlab 19b (MathWorks, Natick, MA, USA).

## 3. Results

From this point on, we will refer to the entire system as Neuromorphic Processing System (NPS), which includes both NAS and SpiNNaker.

### 3.1. Comparison of NPS with spike-based algorithms

In order to evaluate the performance of NPS in detecting bursts of electrophysiological activity, we selected a set of spike-based burst detection algorithms (see section 2.3.2). We then compared the results of VI (i.e., our ground truth) with those of NPS and of the selected spike-based methods. We first computed the number of detected bursts, for which NPS and VI showed no differences, while ISIrank exhibited statistical difference with respect to LogISI, MI and HSMM (Fig. 4, panel A). Given the fact that all the methods produced comparable results, we further investigated the duration of the identified events. Again, no statistical differences were observed between NPS and any of the other detection methods, including VI (Fig. 4, panel B). The only statistical difference was observed between VI and RS, which exhibited the smallest burst duration. In terms of correlation, the burst signals coming from VI and all the other methods showed neither qualitative (Fig. 4, panel C1) nor quantitative (Fig. 4, panel C2, C3) changes in terms of max correlation peak or lag at maximum peak. As expected, NPS showed a larger jitter with respect to the other methods because of the 50ms (non-overlapping) bins. Overall, the NPS results appeared coherent with all the best spike-based burst detectors with an acceptable higher jitter due to the online and 50ms binned procedure.

**Figure 4:**
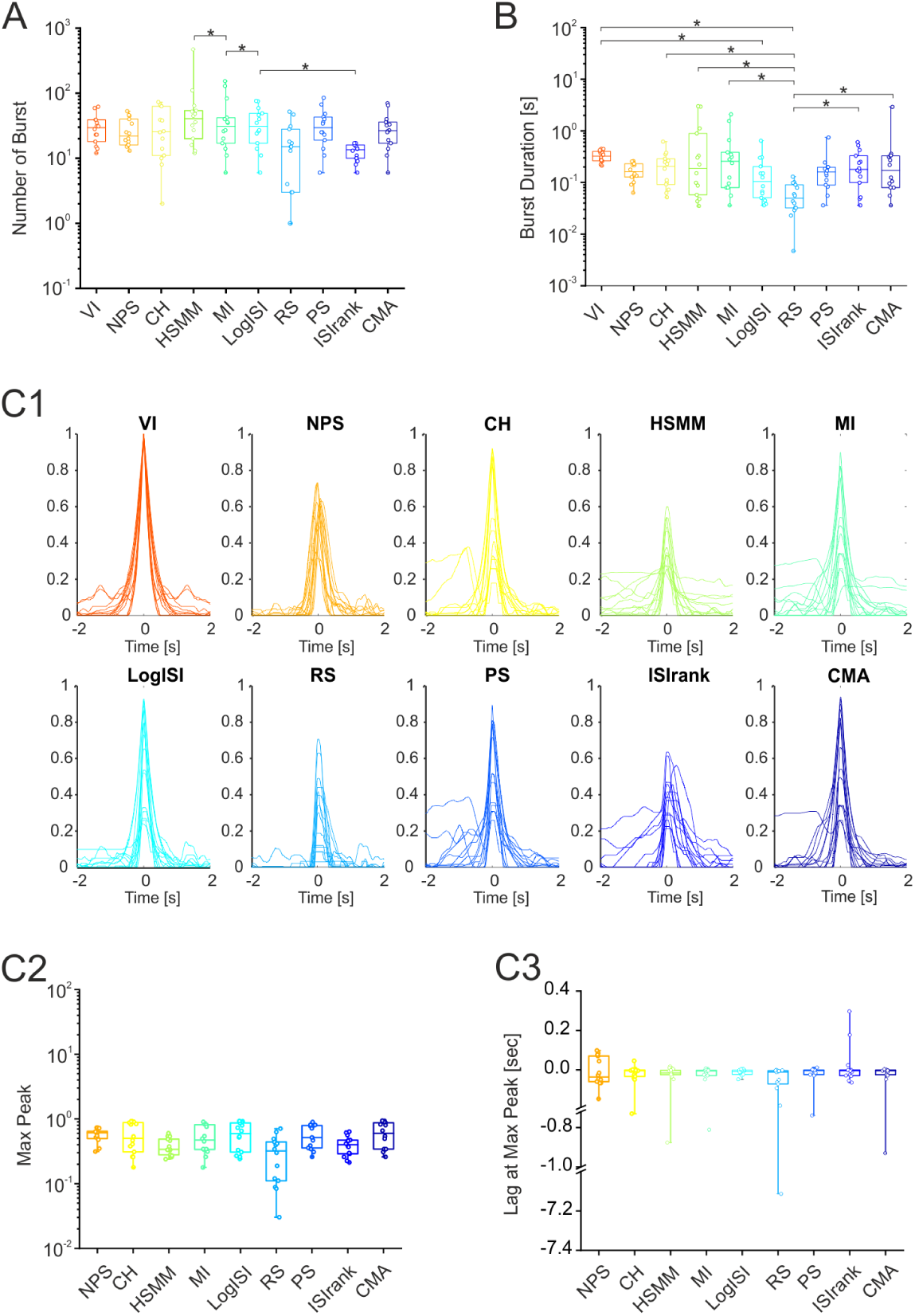
Comparison between NPS, VI and all the spike-based methods. A) Number of burst events detected in 5 minutes for all the 14 recordings belonging to the test set. B) Average burst duration for all the 14 recordings. C1) Qualitative comparison of the cross correlation functions for all methods vs VI (for all the 14 recordings). The top left panel represents the auto-correlation functions of VI. C2) Max Peak values of all the correlation functions reported in panel C1. C3) Lags (expressed in seconds) at the peak of the cross correlation functions. For each box plot, the central line indicates the median and the box limits indicate the 25th and 75th percentiles. Whiskers represent the 5th and the 95th percentiles. Y-axis breaks were done to allow for the visualization of all data points in panel C3. The statistical analyses were carried out using the Kruskal-Wallis test with the Bonferroni post-hoc correction. See Table 2 for the exact p values.

As was already mentioned, most of the current burst detection algorithms rely on spike detection, a process which is prone to errors. Indeed, whenever the high frequency components of the bursts (i.e., the spikes) are not present or are not correctly captured by the detection algorithm, the subsequent burst identification is not reliable. This problem is well-depicted in Fig. 5: three bursting events are clearly visible in the raw and filtered trace; while the first and second bursts are correctly identified by all the spike-based methods (except for RS), the third one is not. Only the NPS is able to capture the burst, which is also correctly identified by the VI. Then, to correctly evaluate the performance of the NPS, we decided to compare it with other raw-based burst detection methods.

**Table 2:**
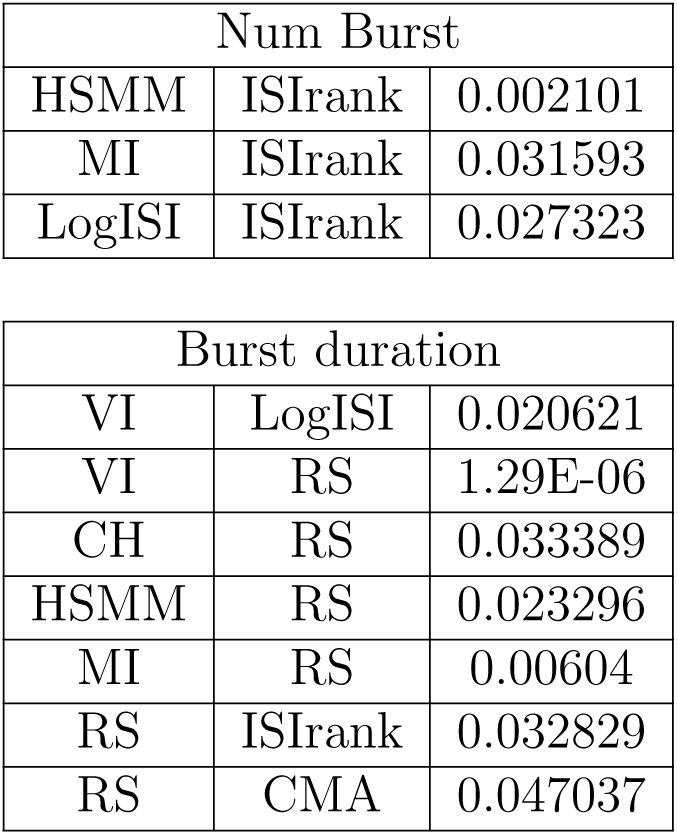
Statistical differences, p*<*0.05, using Kruskal-Wallis test with the Bonferroni post hoc correction. Data referred to Fig. 4.

**Figure 5:**
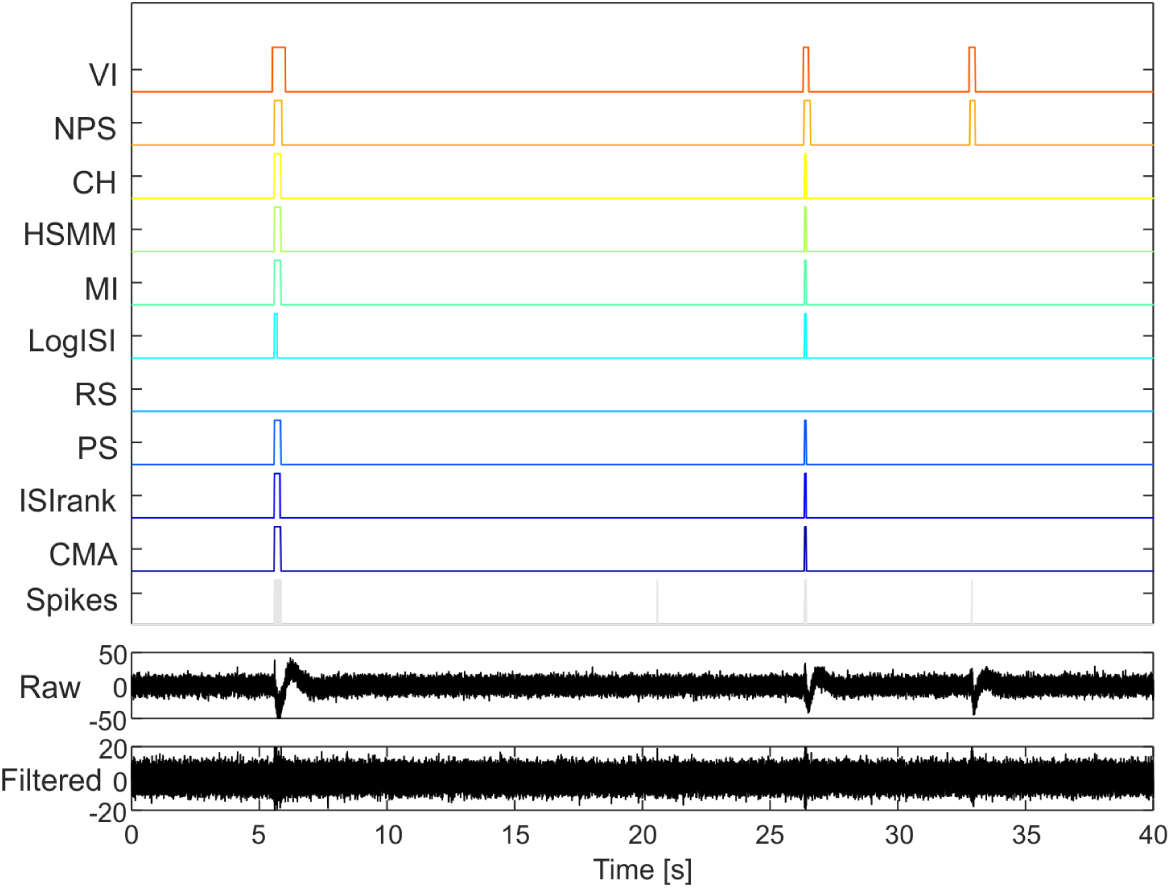
Qualitative comparison between all burst detection methods. Spikes are depicted in grey (all methods, excluding VI and NPS, used spikes to detect bursts). The two signals on the bottom of the figure represent the raw (on which the NPS approach was tested) and the high-pass filtered data (on which we performed the spike detection - using PTSD as described in section 2.3.1).

### 3.2. Comparison of NPS with raw-based algorithms

We tested three burst detection methods based on computing three simple features of the raw signal (i.e., one for each method, namely *max, peak peak, len*) within time windows of 50 ms, as the one used for NPS. Once the training set thresholds were found (see section 2.4), we applied them to the test set recordings. The results in Fig. 6 (panel A) show that only the NPS was able to identify a number of bursts comparable to those of VI. For the three raw-based methods, instead, the number of detected bursts was orders of magnitude higher than the true one. Indeed, the number of false positive bursts was extremely large and, thus, unacceptable. Moreover, the *len* method erroneously counted zero bursts in 8 out of 14 recordings, meaning that such a parameter was not reliable and, therefore, useless for this comparison. We then investigated other parameters, specifically burst duration (Fig. 6, Panel B) and cross correlation peaks and lags (Fig. 6, Panels C1, C2 and C3), Quantitatively, the burst duration (Fig. 6, panel B) of *peak peak* was lower than VI and NPS (p*<*0.05) and *max* showed lower values than NPS (p*<*0.05) while *len* was higher than *peak peak* (p*<*0.05). No statistical differences were found between VI and NPS.

**Figure 6:**
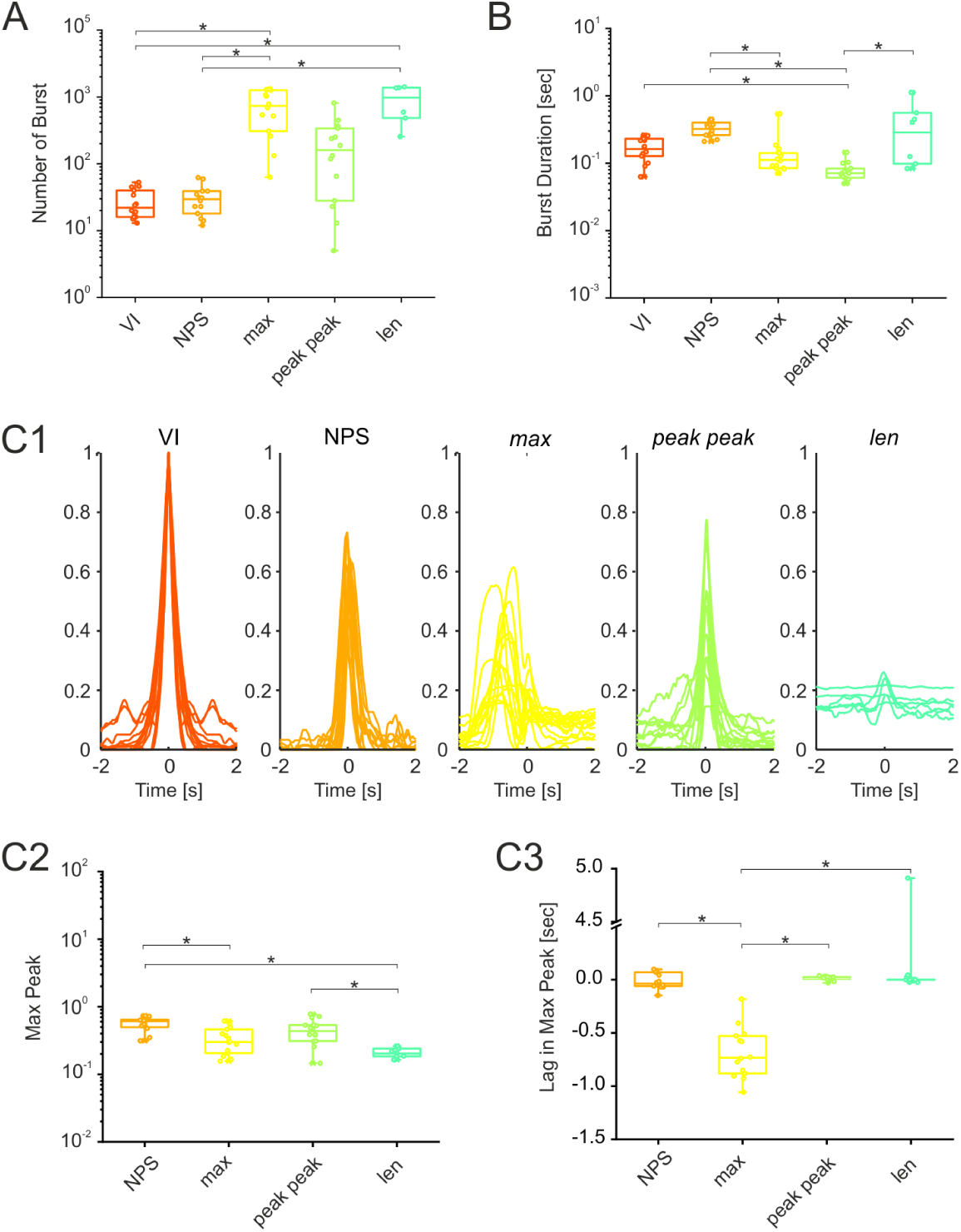
Comparison between NPS, VI and all the raw-based methods. A) Number of burst events detected in 5 minutes for all the 14 recordings belonging to the test set. B) Average burst duration for all the 14 recordings. C1) Qualitative comparison of the cross correlation functions for all methods vs VI (for all the 14 recordings). The top left panel represents the auto-correlation functions of VI. C2) Max Peak of all the correlation functions reported in panel C1. C3) Lags (expressed in seconds) at the peak of the cross correlation functions. For each box plot, the central line indicates the median and the box limits indicate the 25th and 75th percentiles. Whiskers represent the 5th and the 95th percentiles. Y-axis breaks were done to allow for the visualization of all data points in panel C3. The statistical analyses were carried out using the Kruskal-Wallis test with the Bonferroni post-hoc correction. See Table 3 for the exact p values.

Regarding cross correlation, we could observe that the cross-correlation between VI and NPS, *max, peak peak* was qualitatively good and similar to VI’s autocorrelation. Instead, *len* gave again the worst results. Focusing on panels C2 and C3 of Fig. 6, we found that, also in this case, VI and NPS showed no difference. Regarding the Max Peak, the only statistical differences were found between: NPS and *max*, NPS and *len*, and *peak peak* and *len* (p*<*0.05). Regarding the lag, *max* was statistically lower than all the other methods (p*<*0.05), again indicating that it could not be considered as a good alternative to NPS. It is worth highlighting that in 5 out of 14 recordings, the Max Peak of the correlation was higher for *peak peak* than NPS. This does not necessarily suggests that the burst detection in those cases was better for *peak peak* than for NPS. It actually means that, for those cases, VI bursts had a higher overlap with *peak peak* compared to NPS. Indeed, as also suggested by the results on the number of detected bursts (Fig. 6, panel A), the amount of false positives is huge, as qualitatively reported in Fig. 7. Therefore, the overall performance is much higher for NPS when compared to *max, peak peak* and *len*.

**Table 3:** Statistical differences, p*<*0.05, using Kruskal-Wallis test with the Bonferroni post hoc correction. Data referred to Fig. 6.

| Num Bursts |  |  |
| --- | --- | --- |
| VI | *max* | 1.52E-05 |
| VI | *len* | 0.0002977893 |
| NPS | *max* | 4.24E-05 |
| NPS | *len* | 0.0005998977542 |

| Burst duration |  |  |
| --- | --- | --- |
| VI | *peak peak* | 0.01123345496 |
| NPS | *max* | 0.003838358311 |
| NPS | *peak peak* | 7.85E-08 |
| *peak peak* | *len* | 9.32E-03 |

| Max Peak |  |  |
| --- | --- | --- |
| NPS | *max* | 0.005132954712 |
| NPS | *len* | 0.0003261603306 |
| *peak peak* | *len* | 0.03145223724 |

| Lag |  |  |
| --- | --- | --- |
| NPS | *max* | 0.0005797489213 |
| *max* | *peak peak* | 1.06E-06 |
| *max* | *len* | 3.61E-05 |

**Figure 7:**
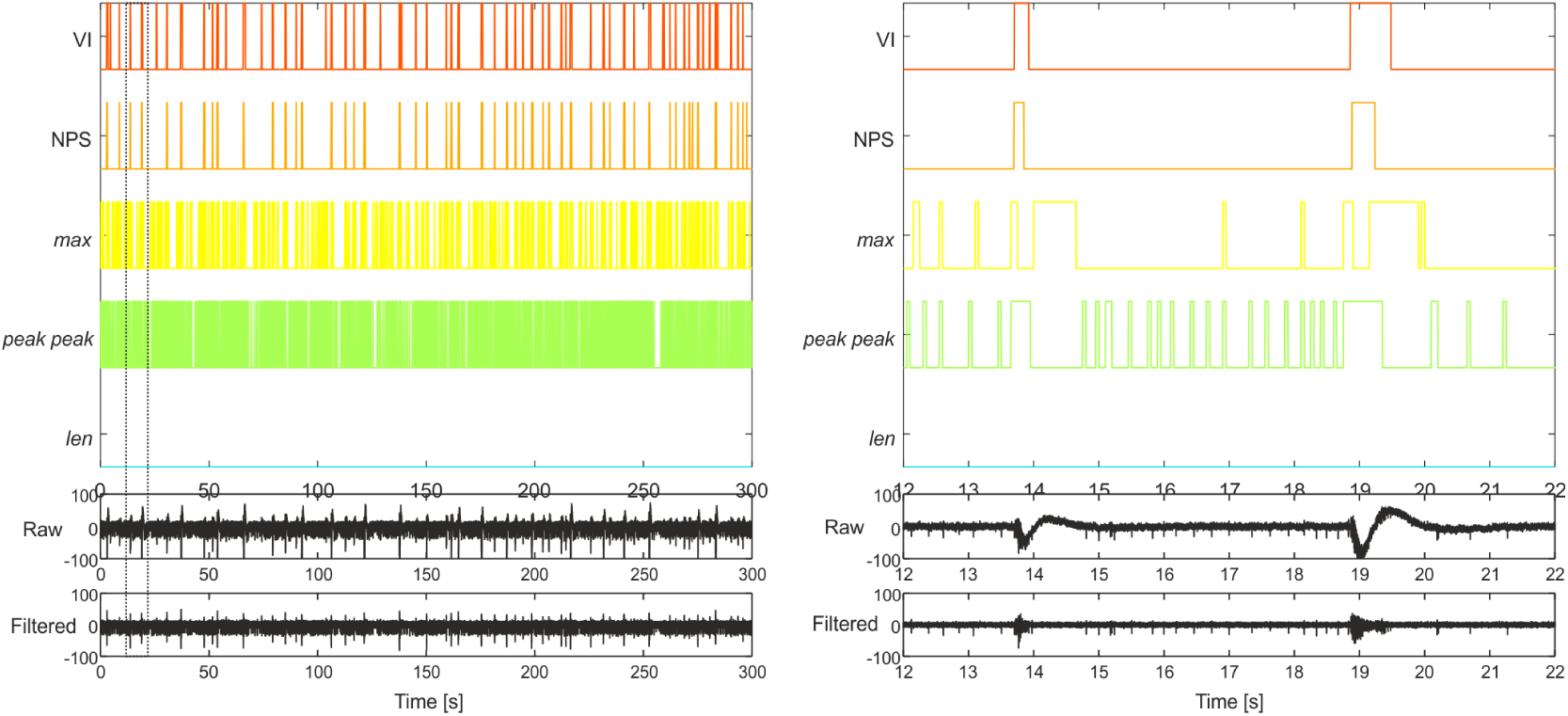
Qualitative comparison between all raw-based burst detection methods. On the left, 300s recording from one of the test set electrodes. The two signals on the bottom of the figure represent the raw (on which the NPS approach was tested) and the high-pass filtered data. The dotted rectangle shows the 10s detail reported on the right. In this case, the *len* method was not able to identify any burst event.

**Figure 8:**
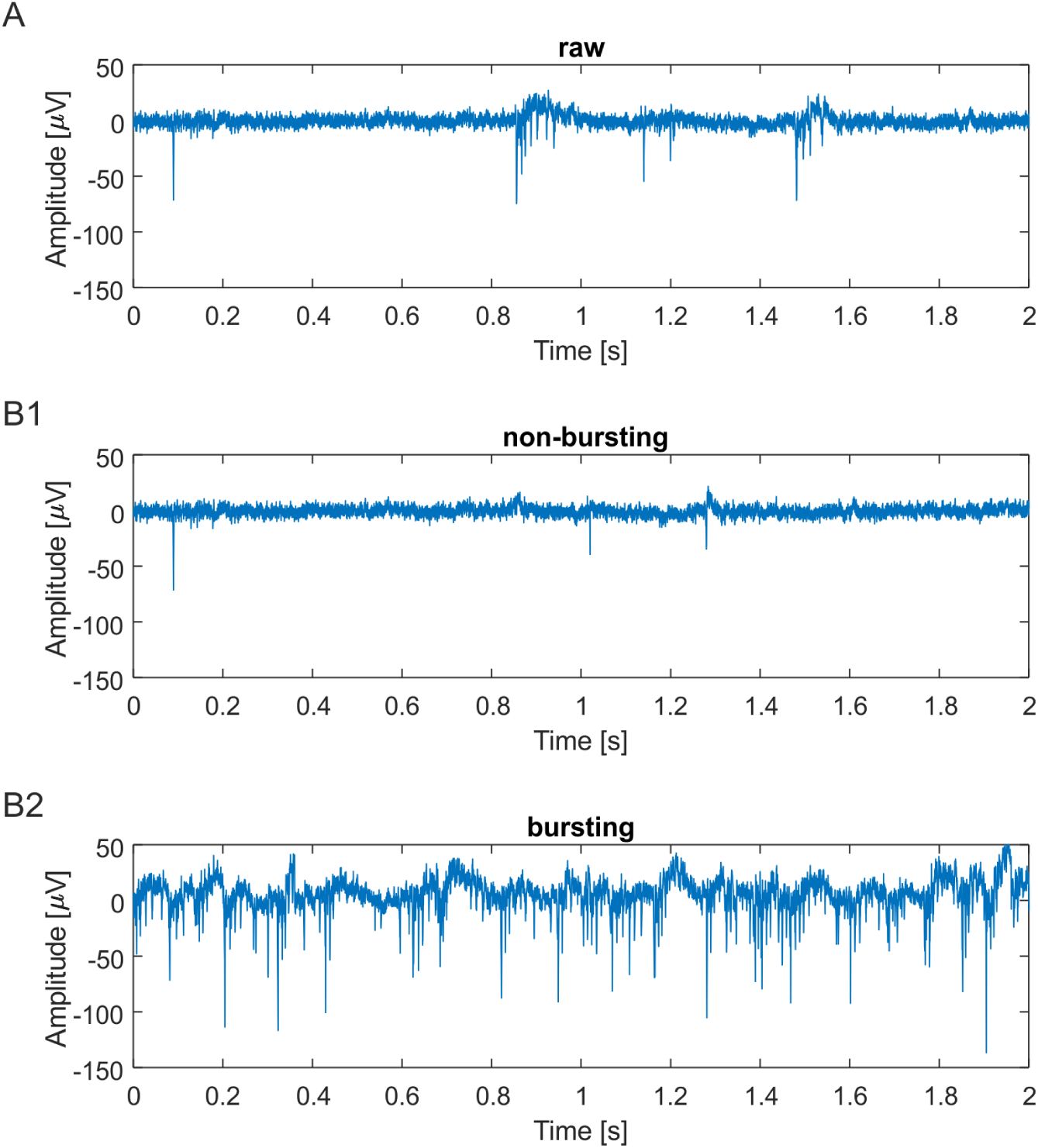
Comparison of training classes. A) First 2 seconds of raw signal from the training set. B) First two seconds of “non-bursting” training set. C) First two seconds of “bursting” activity.

**Figure 9:**
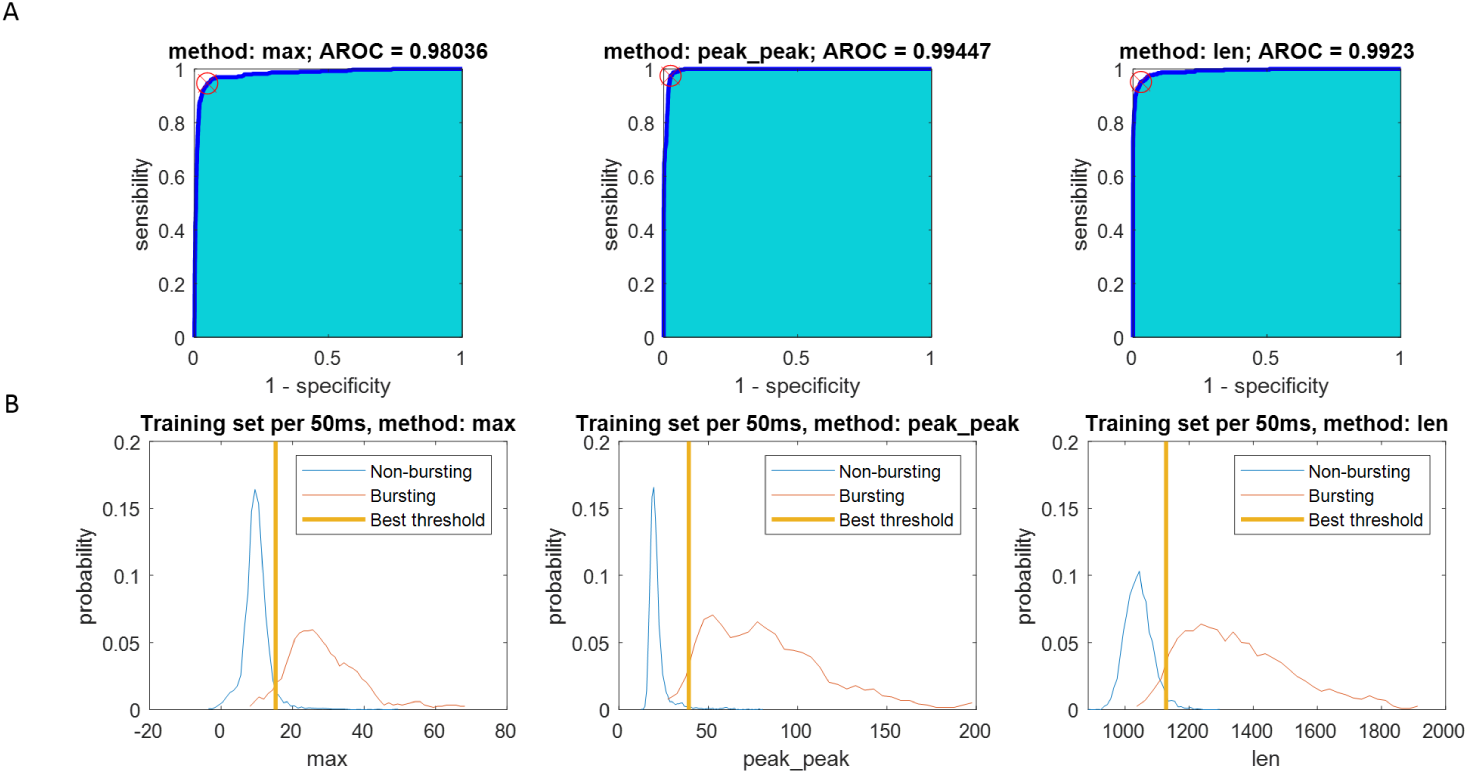
Training set thresholds. A) ROC curves with superimposed red cross and circle representing the best threshold (i.e. the one that minimizes the distance from the top-left corner) B) histograms of bursting (red) and non-bursting (blue) windows for each method (max, *peak peak* and *len*). The yellow vertical line represents the best threshold.

## 4. Discussion and conclusion

In biological neural network recordings, the correct identification of burst events is crucial in many scenarios, ranging from basic neuroscience to biomedical applications [51, 16]. Burst events are characterized by both a high-pass filtered component (highly packed spikes) and a low frequency component (local field potentials LFP) due to the synaptic currents.

Although several burst detection methods have been proposed in the literature, none of them have been widely adopted to date [16].

Here we introduce an innovative burst detection technique based on a neuromorphic, data-driven, online approach. We used the Neuromorphic Auditory Sensor (NAS), which was originally designed to work with audio signals and convert them into spiking events with a frequency decomposition approach. For this work we used the raw data (containing the full spectrum) coming from electrophysiological recordings of neuronal cultures plated over Micro Electrode Arrays. The NAS converted the raw activity of neurons in an event-based signal which was then fed into the neuromorphic SpiN-Naker board, previously trained to detect bursting events. For the sake of simplicity, the complete burst detection process was named NPS. In order to evaluate its performances, we initially compared it with 8 conventional (i.e., spike-based) methods widely used in the literature [16] and a visual inspection (VI). Conventional burst detection methods rely on a previous spike detection step. Unfortunately, spike detection can be biased by the background noise [52, 60]. In case of chronic recordings, the high-frequency (*>*300Hz) component of the signal can be degraded by the micro-motion of the electrodes and inflammatory tissue response [61], thus severely affecting the detection performances. Conversely, low frequency components, which are usually overlooked, can be phase-locked to the bursting activity [62, 63] and more stable in time [64, 65]. Considering the multi-frequency content of a neural signal, NAS is able to analyze both the low and the high frequency components and, therefore, can easily generalize across different recordings. Specifically, for long-term in vitro [66, 13, 67] and in vivo [64] studies, where the signal-to-noise ratio can change significantly, our approach can highlight meaningful events regardless of the environmental conditions. Moreover, the ability of our approach to detect low frequency events can be used to extract network-wide events (such as Network Burst [68]) from a single recording site, thus providing a larger overview of the background activity of the neuronal network without the need of multiple recording sites.

Another drawback of spike-based burst detection methods is the fact that they are generally implemented offline. Among the conventional algorithms considered for this study, only few can be redesigned to work online, such as CH [58], while others need to know the entire distribution of spike intervals in advance, such as LogISI [56], and therefore are hardly convertible to work online. On the contrary, NPS has been designed as a native online method, thus making it a perfect candidate for neuroengineering applications and devices based on closed-loop approaches [69, 70, 71].

In terms of quantitative results, it is worth mentioning that our NPS approach was similar in terms of the number of detected events, mean burst duration and correlation to VI and most of the tested spike-based burst detectors. The main difference with spike-based methods was related to the higher lags with respect to the ground truth due to the 50ms-long time windows used, which reduced the precision of the detection time. To reduce this lag we could decrease the time window used by NPS. At the same time, reducing the time window would result in a possible fragmentation of the burst events due to smaller silent periods or with less activity within the same event. Therefore, an accurate choice should be made to obtain the best from this trade-off.

Besides spike-based approaches, as a second step, we compared our method with a set of raw-based methods trained and tested on the same dataset. The results show that a simple feature extracted from 50ms-long time windows is not enough to set a threshold which can correctly discriminate between bursting and non-bursting activities on the test set. In general, other methods, including machine-learning and deep-learning techniques, could be implemented to perform the same task. However, an event-based approach as the one implemented here has a competitive advantage over traditional methods, which is the reduced power consumption and computational latency [72]. An additional point to mention is that in traditional audio/video processing systems, the information is processed periodically following a sampling rate. However, in these systems, not all samples contain relevant information (e.g., when no audio is obtained in the input, or when two consecutive frames from a camera are the same), but the system still has to process them. On the other hand, Neuromorphic systems and sensors only process the input when there is relevant information. In this case, when no audio signal is received by the NPS, the system does not perform any computation, staying idle and, thus, reducing power consumption. Moreover, the hardware implementation of this system is a natural improvement when compared to the software, offline methods used as comparison. Having a hardware system already available and the possibility to use electrophysiological recordings as input, constitutes an advantage with respect to software-based methods. Recently published neuromorphic-based studies, exploited the potential application of SNNs in combination with their biological counterparts [73, 74]. Others have started using SNNs for pattern recognition in biomedical applications on humans. A recent work ([75]) exploited a SNN approach to discriminate electromyography (EMG) signals. These were also used in a previous study by Peng et al. [76] for a 6-class recognition problem of hands motion with a 1k neurons SNN. Others have used SNNs for classifying Electroencephalography (EEG) signals [77, 78].

Our work is along the same line and all the mentioned peculiar features of our neuromorphic approach suggest that a neuroprosthetic application can be the natural, future evolution of this work.

## Acknowledgements

Preliminary results were obtained at the Capo Caccia Cognitive Neuromorphic Engineering Workshop (Apr 24 - May 06, 2017, Capo Caccia, Italy) by Juan P. Dominguez-Morales, Stefano Buccelli, Daniel Gutierrez-Galan and Ilaria Colombi: all the authors would like to thank all the organizers of the Workshop.

This work was supported by the Spanish grant (with support from the European Regional Development Fund) COFNET (TEC2016-77785-P). The work of Juan P. Dominguez-Morales was supported by a Formación de Personal Universitario Scholarship from the Spanish Ministry of Education, Culture and Sport. The work of Daniel Gutierrez-Galan was supported by a Formación de Personal Investigador Scholarship from the Spanish Ministry of Education, Culture and Sport.

## 5. Supplementary

## Footnotes

1 http://neuralensemble.org/docs/PyNN/reference/plasticitymodels.html

